# *Candida albicans* biofilms are generally devoid of persister cells

**DOI:** 10.1101/420596

**Authors:** Iryna Denega, Christophe d’Enfert, Sophie Bachellier-Bassi

## Abstract

*Candida albicans* is known for its ability to form biofilms – communities of microorganisms embedded in an extracellular matrix developing on different surfaces. Biofilms are highly tolerant to antifungal therapy. This phenomenon has been partially explained by the appearance of so-called persister cells, phenotypic variants of wild-type cells, capable of surviving very high concentrations of antimicrobial agents. Persister cells in *C. albicans* were found exceptionally in biofilms while none were detected in planktonic cultures of this fungus. Yet, this topic remains controversial as others could not observe persister cells in biofilms formed by the *C. albicans* SC5314 laboratory strain. Due to ambiguous data in the literature, this work aimed to reevaluate the presence of persister cells in *C. albicans* biofilms. We demonstrated that isolation of *C. albicans* “persister cells” as described previously was likely to be the result of survival of biofilm cells that were not reached by the antifungal. We tested biofilms of SC5314 and its derivatives, as well as 95 clinical isolates, using an improved protocol, demonstrating that persister cells are not a characteristic trait of *C. albicans* biofilms. Although some clinical isolates are able to yield survivors upon the antifungal treatment of biofilms, this phenomenon is rather stochastic and inconsistent.

## INTRODUCTION

The yeast *Candida albicans* is a commensal of humans but also one of the most prevalent fungal pathogens, responsible for superficial infections as well as life-threatening systemic infections (1). *C. albicans* is recognized for its ability to form biofilms that are most frequently associated with nosocomial infections, particularly in immunocompromised patients.

*C. albicans* biofilms are communities of microorganisms with a complex structure composed of different cell types embedded in an extracellular matrix (2–4). They develop on different types of surfaces, either living or inert, and are characterized by their high tolerance to antifungals. The latter can result from the properties of the extracellular matrix that can serve as a trap for drug molecules (5–7). An additional source of antifungal tolerance has been proposed to result from the occurrence in biofilms of so-called persister cells, a subpopulation of phenotypic variants of wild-type cells, capable of surviving concentrations of antimicrobial agents well above the Minimal Inhibitory Concentration (MIC) (8). Persister cells are genetically identical to other biofilm cells. Upon removal of the antimicrobial agent they give rise to a new population comprised of the majority of susceptible cells and a new small subpopulation of persisters. Thus, persistence is a non-inherited trait (9–11).

In the clinical setting, persisters are usually associated with relapse of infections and with the development of chronic infections. For bacterial persisters, several mechanisms and pathways involved in their development have been described (12). In 2006, LaFleur et al. have presented the first report of persister cells in biofilms of *C. albicans*, which could contribute to biofilm tolerance to antifungals (8). In their paper the authors have reported that *C. albicans* exhibit a biphasic killing curve, when exposed to the antifungals such as amphothericin B (AMB), chlorhexidine or the combination of both. This phenomenon is explained by the presence of a multidrug-tolerant subpopulation of persister cells within a biofilm. Notably, the experiments for this study were performed using in vitro biofilm model of *C. albicans*, developed in polystyrene 96-well plates. Following this work and relying on the protocol for persister cells isolation described therein (8), persister cells in *C. albicans* biofilms were described by a few other groups (13–15). However, later work by the Douglas group showed that not all Candida species and strains were able to form persister cells in laboratory-grown biofilms (16). This was in particular the case for *C. albicans* strain SC5314 (17), the parental strain of almost all *C. albicans* strains used for functional genomics and molecular genetics studies. Unlike in the previously mentioned papers (8, 13–15), the protocol Al Dhaheri and Douglas (16) used for persisters isolation involved growing biofilms on silicone discs followed by their immersion into an antifungal solution. As the topic of *C. albicans* persister cells remains controversial, the main objective of this work was to reevaluate their occurrence in *C. albicans* biofilms.

## METHODS

### Strains and growth conditions

In this study we used 3 reference strains (listed in Table 1) and a set of 95 clinical isolates (Table S1).

**Table. 1.** *C. albicans* strains used in this study

| STRAIN | GENOTYPE | REFERENCE |
| --- | --- | --- |
| SC5314 |  | (17) |
| CEC369 | *ura3::λimm434/ura3::λimm434 ARG4/arg4::hisG*<br>*HIS1/his1Δ::hisG RPS1/RPS1::Clp10* | (18) |
| CEC4664 | *ura3Δ::λimm434/ura3Δ::λimm434*<br>*iro1Δ::λimm434/iro1Δ::λimm434 ADH1/adh1::P<sub>TDH3</sub>-*<br>*carTA::SAT1 arg4Δ/ARG4 his1Δ::hisG/HIS1*<br>*RPS1/RPS1::Clp10* | Lab's<br>collection |

Yeast precultures were grown overnight in YPD (1% yeast extract, 2% peptone, 2% glucose) with shaking at 30^°^C.

Biofilms were grown either in RPMI 1640 medium with L-glutamine (buffered with 50 mM HEPES), as described in (8) and (18), or in GHAUM medium (SD supplemented with 2% glucose and 1 mg/mL histidine, 1 mg/mL arginine, 0.02 mg/mL uridine and 2 mg/mL methionine (19)).

### Biofilm growth and persister cells isolation

To assess persister cell appearance in biofilms we used two protocols adapted either from (8) or (13). The first protocol uses 96-well plates and the biofilms are grown in RPMI. In the second protocol the biofilms are grown in 24-well plates but using GHAUM medium instead of YNB.

#### Biofilm growth

Overnight cultures were washed in sterile 1x PBS and diluted in the corresponding medium to OD_600_ 0.3. Either 100 µL or 1 mL of cells in the 96-well plate or the 24-well plate, respectively, were allowed to adhere for 1.5 h without agitation. The non-adhered cells were then washed with 1X PBS, the same volume of fresh medium was added, plates were covered with a breathable seal and biofilms were allowed to form for 24 h at 37^°^C with agitation (110 rpm). At this point the media were changed and biofilms were allowed to grow for 24 more hours.

#### Antifungal treatment

Old media were carefully aspirated, without disrupting the biofilm structure. Biofilms were washed once with either 100 µL or 1 mL of 1x PBS, respectively, and treated with a 100 µg/mL AMB solution in either RPMI or GHAUM for 24 hours at 37^°^C, statically. AMB solutions were prepared from a 8 mg/mL stock in DMSO, so that the final concentration of DMSO in a working solution did not exceed 1.25%. For control biofilms, corresponding amount of DMSO was added to the medium instead of the antifungal solution.

This step was either performed using the same volumes of antifungal solution as for biofilm growth as described in (8) and (13). or increasing the volume of antifungal to fill the well up to the top (350 µL or 3 mL for 96- and 24-well plates, respectively). Clinical isolates were first treated with 64 µg/mL AMB solution. Strains giving rise to colonies were then tested 5 times with 100 µg/mL AMB.

#### Plating

Upon 24 hours of antifungal treatment, AMB solution was aspirated and biofilms were washed twice with 1X PBS prior to plating on YPD-agar plates. Biofilms were resuspended in 1x PBS/0.05% Tween-20. For the AMB-treated samples, the whole biofilms were plated. For control biofilms, serial dilutions were performed to allow CFU counting. CFU were counted after incubating the plates at 30^°^C for 48 h.

## RESULTS AND DISCUSSION

In this work, we aimed to study the occurrence of persister cells in *C. albicans* biofilms. We applied the protocol published by LaFleur and colleagues, growing the biofilms in RPMI and in a 96-well plate format (8). We set up the protocol with 3 *C. albicans* prototroph strains, namely SC5314, CEC369 and CEC4664 - prototroph derivatives of BWP17 and SN76, respectively. BWP17 (20) and SN76 (21) are independent auxotroph derivatives of SC5314 and have been rendered prototroph with sequential transformation events.

We encountered a technical problem at the biofilm recovery step, usually performed by scraping the cells in 1x PBS and vortexing prior to plating (8, 13, 15, 22). In our hands, the cells could not be properly resuspended and plated, as clumps of the biofilms would usually remain stranded inside the tips. Consequently, the CFU numbers obtained were highly variable for all samples, making any further analysis and comparison impossible (data not shown).

We decided to test alternative approaches to circumvent the stickiness of biofilms. Resuspending cells in 20% glycerol/1X PBS for plating helped reducing stickiness, but did not improve consistency (data not shown). We hypothesized that EDTA might reduce adherence of biofilms by binding bivalent cations that are required for the activity of cell surface adhesins (23). Thus, we attempted applying 20% glycerol with a range of EDTA concentrations (0, 50, 100 mM) for plating. 100 µL of EDTA solutions of different concentrations were added to biofilms and left for 10 minutes at room temperature prior to biofilm disruption by scraping and vortexing. None of the applied EDTA solutions allowed abolishing stickiness. Additionally, colonies growing on YPD-agar exhibited a wrinkled morphology, most probably linked to the toxicity of ED TA (24). Finally, we tried adding Tween-20 (0.05%) to PBS. Tween-20 eradicated the problems of stickiness and poor disruption and improved recovery of cells from the biofilms (Fig. 1). The effect on cell viability was tested using a planktonic culture of SC5314 that was washed and plated on YPD-agar using PBS and PBS-Tween-20 solutions. No impact on viability was observed (data not shown). Thus, in the experiments described below, biofilms were resuspended in a 0.05% Tween-20/1X PBS solution.

**Fig. 1.**
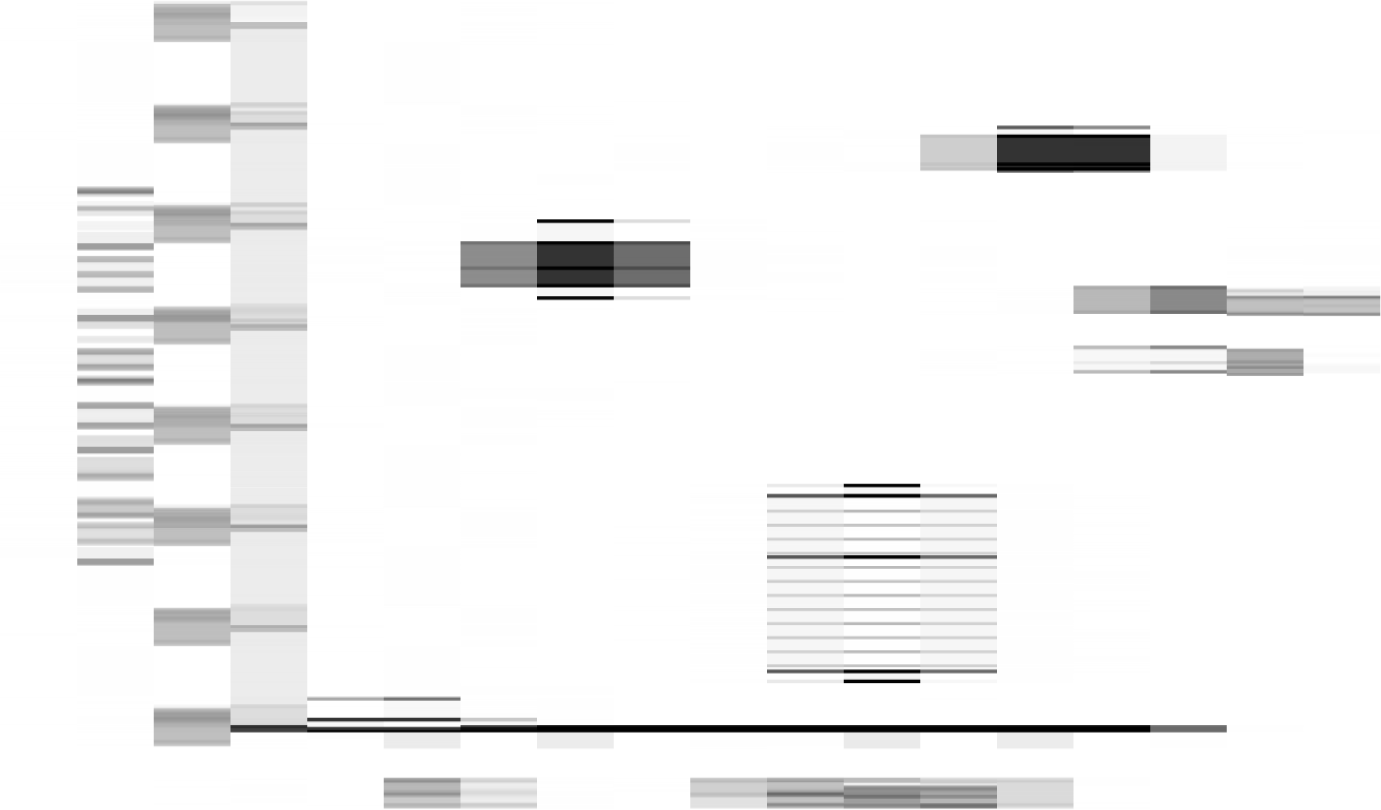
Effect of Tween 20 on the recovery of CFUs from *C. albicans* SC5314 biofilms. *C. albicans* SC5314 was allowed to form biofilms in 100 μL RPMI in a 96-well plate according to the protocol adapted from (8). Error bars: standard deviation (SD) of 6 biological replicates generated from 2 independent experiments.

However, even after this modification, the ratio of cells that survived AMB treatment was still inconsistent between repeats. According to Lafleur and colleagues the ratios of *C. albicans* persister cells in biofilms vary from 0.1% to 2% for different strains, notably from 0.05 to 0.1% for strain CAI4 – a derivative of *C. albicans* SC5314 (8). Our values hardly ever exceeded 0.01% persisters per biofilm, even after improving the recovery protocol, thus bordering with statistical error. We reasoned that increasing the surface of a biofilm and changing the growth media could improve persister yields and decided to test the protocol described in (13), applying the modifications that were mentioned previously. However, the problem of inconsistency and low ratios of persisters remained (Fig. 2).

**Fig. 2.**
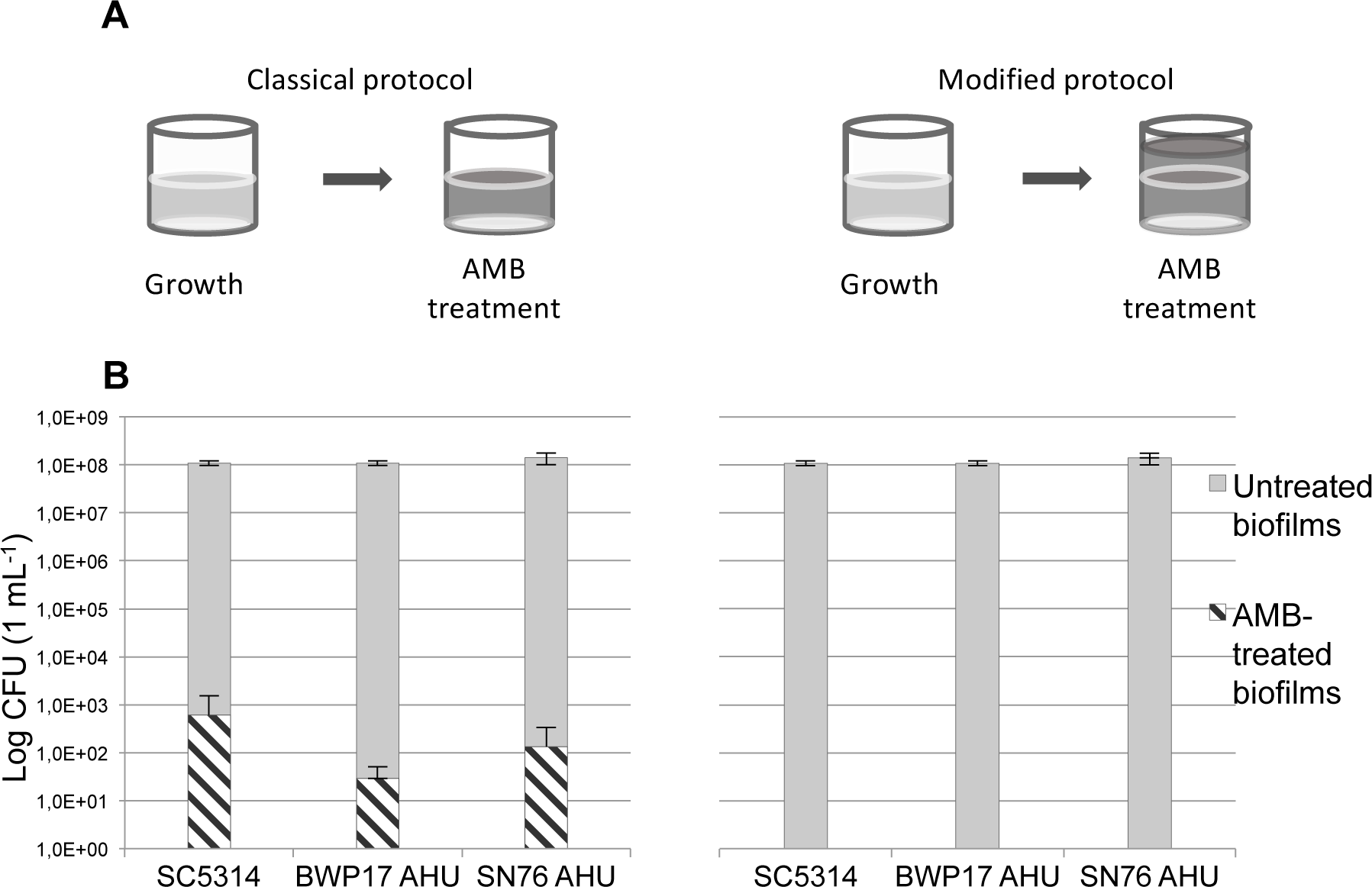
Schemes of the protocols (A) and levels of persisters (B) obtained from biofilms grown using modified protocol from (13). Biofilms were grown in 1 mL of GHAUM medium in 24-well plates before application of either 1 mL of AMB solution (on the left) or 3 mL of AMB solution (on the right). Ratios of surviving cells are as follow: SC5314 – 5.6*10^-4^%, CEC369 – 2.6*10^-5^%, CEC4664 – 9.4*10^-5^%. Error bars: SD of 6 biological replicates generated from 2 independent experiments.

In all protocols described previously, the volumes of the media and solutions used for biofilm growth, washing, and AMB treatment were identical. Upon a careful observation, we noticed that *C. albicans* cells form a dense rim at the border of the air and liquid phases, as a result of agitation during growth. Treating a biofilm with the exact same volume of antifungal and growth medium in static conditions thus could result in cells from the rim escaping treatment. We decided to increase the volume of the applied antifungal solution (filling wells to the top) and, to our surprise, this change in the protocol led to a complete eradication of persisters for the laboratory strain SC5314 and its derivatives. Reproducibly, we did not get any persisters after applying this change for all strains for both RPMI- and GHAUM-grown biofilms. Thus, the volume of the antifungal applied in the original protocols for persister isolation was skewing the results. Increasing the volume of antifungal eliminated this bias, resulting in a complete eradication of any survivors after the antifungal treatment.

In our work we used a modified protocol for persister cells isolation with a starting cell suspension of OD_600_ 0.3 used for biofilm growth instead of 0.1 as described in the original protocols (8, 13). To assess the impact of the initial cell number used for seeding biofilms on persister cells’ appearance, we tested our protocol for SC5314 using cell suspensions of OD_600_ 0.1, 0.3 and 0.5 for seeding. Regardless of the initial biomass, persister cells did not form in SC5314 biofilms grown either in RPMI or GHAUM (data not shown).

These results made us question the very existence of persister cells in *C. albicans* biofilms. Previously, Al-Dhaheri and Douglas showed that not all strains of *C. albicans* can form persister cells (16). Particularly, in their hands, SC5314 biofilms lost all viability after exposure to 30 μg/mL AMB. However, biofilms of another clinical isolate, GDH2346, appeared to contain a small proportion (0.01%) of cells that survived 100 μg/mL AMB treatment. These authors used a different *in vitro* model for assessing persistence, as they grew biofilms on silicone disks that were transferred to a new well filled with an antifungal solution. This prevented an escape of any cells from the antifungal treatment. Thus, our modified protocol for treatment of biofilms formed in 96-well or 24-well plates corroborated the results obtained by the Douglas group for *C. albicans* strain SC5314 (16).

Since the clinical isolate GDH2346 could give rise to survivors (16), we could not exclude that persisters could emerge in biofilms of different *C. albicans* isolates. Additionally in 2010, LaFleur and colleagues isolated and described *C. albicans* strains from patients with long-term oral infection, that gave yield to increased levels of persisters (up to 8.9%) (23). These were called *hip*-mutants, by analogy with the high persister strains previously described for bacteria (26, 27). Although *hip*-mutants were identified using a protocol that showed limitations in our hands, we hypothesized that some *C. albicans* clinical isolates could generally be more prone to form persisters than others (namely SC5314). To test this assumption, we tested 96 clinical isolates (Table S1) for their ability to form biofilms and the occurrence of persister cells following AMB treatment. In a first round of experiments, biofilms were treated with a 64 µg/mL AMB solution. Only 38 isolates (39.6%) displayed survivors (notably, never exceeding a rate of 0.02%). According to the generally accepted concept of persistence (9), the frequency of persisters’ appearance is independent of the increase in antibiotic concentration. Thus in a second round of experiments, biofilms were developed for these 38 isolates and treated with a 100 µg/mL AMB solution. Notably, only 7 isolates out of these 38 displayed survivors when grown with 100 µg/mL AMB (CEC3668, CEC4514, CEC4525, CEC3554, CEC3634, CEC3669, CEC4521). These 7 strains, together with 4 other isolates randomly picked in the remaining 31 strains (CEC4512, CEC3706, CEC712, CEC3708), were tested five more times with 100 µg/mL of AMB. In most cases these strains did not yield persister cells (Fig. 3); however, 6 strains gave rise to survivors in one (CEC4525, CEC3634, CEC3669) or two (CEC5414, CEC3554, CEC4521) of the experiments (Fig. 3), which could be explained either by the stochastic nature of persistence as a phenomenon or by technical errors during the experiment.

**Fig. 3.**
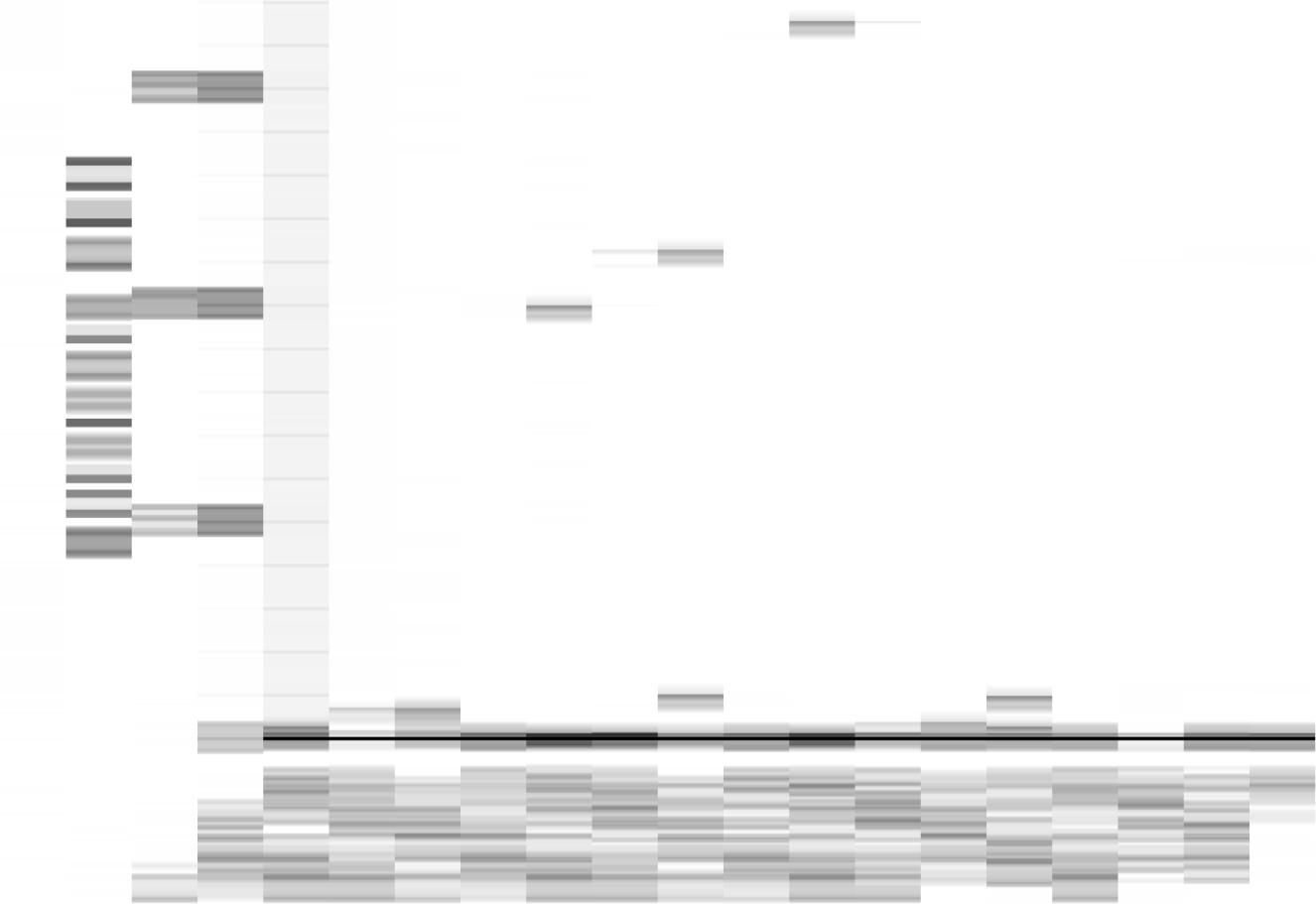
Analysis of persister cell formation in 11 clinical isolates. Biofilms were grown in 1 mL of GHAUM medium in 24-well plates, and treated with 1 mL of AMB solution (modified protocol from (13)). The values obtained from 5 biofilms were used to draw the graph.

## CONCLUSION

Since 1944, when Bigger first described persister cells in Staphylococcus (28), many advances have been made in exploring this phenomenon, especially in bacteria. It is known that microbial cultures growing *in vivo* can sometimes be very difficult to eradicate completely by an antibiotic treatment, causing relapses or development of chronic infections in patients. A small pool of cells with the same genotype as the rest of the population but differing in their ability to tolerate stress – including drug treatment – provides a form of insurance to the population from an evolutionary point of view.

The phenomenon of persistence has been described not only for bacteria, but other types of pathogens, and it has been proposed that persister cells significantly contributed to the recalcitrance of *C. albicans* biofilms to antifungal treatments (29–31).

*C. albicans* persister cells were first described in 2006 (8), and since then just a handful of reports, sometimes contradictory, have been presented. In our study, we explored standard protocols to obtain persisters, and showed that their proportion in biofilms formed by different *C. albicans* strains has been overestimated. Our results show that the detected “persister cells” were likely the result of survival of cells that were not reached by the antifungal. Notably, Al-Dhaheri and Douglas (16) were able to detect some persisters in biofilms of a clinical isolate, but the ratio they obtained was much lower (0.01%) than the numbers published by other authors (8, 13).

Although some of the tested clinical isolates of our study were occasionally able to yield survivors after the treatment of biofilms with AMB, this phenomenon was rather inconsistent, pointing either to the stochastic nature of persistence itself, or another skew in the protocol while carrying out particular experiments.

At that point we cannot completely exclude the possibility of persistence in all existing *C. albicans* strains, though with our protocol we managed to disprove their presence for 91 analysed strains out of 98.

## ACKNOWLEDGEMENTS

Iryna Denega is part of the Pasteur - Paris University (PPU) International PhD Program. This project has received funding from the European Union’s Horizon 2020 research and innovation programme under the Marie Sklodowska-Curie grant agreement No 665807, and from the Institut Carnot Pasteur Microbes & Santé. This work has been supported by grants from the French Government’s Investissement d’Avenir program (Laboratoire d’Excellence Integrative Biology of Emerging Infectious Diseases, ANR-10-LABX-62-IBEID) to C.d’E.

